# Transcriptomic Analysis of 3D Vasculature-On-A-Chip Reveals Paracrine Factors Affecting Vasculature Growth and Maturation

**DOI:** 10.1101/2021.11.21.469415

**Authors:** Sin Yen Tan, Qiuyu Jing, Ziuwin Leung, Ying Xu, Angela R Wu

## Abstract

In vitro models of vasculature are of great importance for modelling vascular physiology and pathology. However, there is usually a lack of proper spatial patterning of interacting heterotypic cells in conventional vasculature dish models, which might confound results between contact and non-contact interactions. We use a microfluidic platform with structurally defined separation between human microvasculature and fibroblasts to probe their dynamic, paracrine interactions. We also develop a novel, versatile technique to retrieve cells embedded in extracellular matrix from the microfluidic device for downstream transcriptomic analysis, and uncover growth factor and cytokine expression profiles associated with improved vasculature growth. Paired receptor-ligand analysis further reveals paracrine signaling molecules that could be supplemented into the medium for vasculatures models where fibroblast co-culture is undesirable or infeasible. These findings also provide deeper insights into the molecular cues for more physiologically relevant vascular mimicry and vascularized organoid model for clinical applications such as drug screening and disease modeling.

## Introduction

At present, there is enormous clinical demand for *in vitro* vasculature models that recapitulate the physiological as well as pathological human blood vessels for a wide range of applications such as drug testing, disease modeling and organoid vascularization. Although much progress has been made to build more physiologically relevant *in vitro* vasculature models, including the use of co-culture systems with other cell types, conventional models generally lack spatial arrangement of those heterotypic cell types in terms of anatomical position and distance. As a result of this lack of appropriate methods to accurately mimic natural vasculature, the paracrine interactions between endothelial cells and their corresponding supporting cells such as fibroblasts have only been studied using models with critical limitations: either using 2D models that lack 3D extracellular matrix (ECM) and do not reflect the cell microenvironment; or using 3D dish models with heterotypic cells in direct contact with each other that misrepresent the physiological condition; or using animal and chimeric models that do not accurately reflect human biology.

An ideal *in vitro* vascular model should fully recreate the complexity of native blood vessel environment while providing simplicity for the operation of the culture system^1^. Microfluidic technology is particularly well suited for this mimicry due to its ability to precisely control both biochemical and mechanical factors regulating the microvasculature environment. Among the distinctive features provided by microfluidic platforms that cannot be achieved in traditional cell culture systems, is a 3D cellular microenvironment that can be customized to create specific spatial compartmentalization of cocultures of multiple cell types, biochemical gradients, and mechanical stimuli such as perfusion flow and shear stress. Most importantly and uniquely, microfluidic platforms enable active fluid perfusion through the vasculature with the opening of the perfusable lumens. In comparison to *in vivo* animal vasculature models such as mice or zebrafish, microfluidic vascularization strategies enable experimentation on a physiologically relevant all-human system, while also minimizing the use of animals, as well as avoiding cross-species contamination that could confound study results^2^.

Of all the supporting cells, fibroblasts have been given much attention since they generally co-exist with endothelial cells *in vivo*. Under normal physiological condition, fibroblasts regulate and maintain ECM homeostasis while secreting essential growth factors and chemokines. In response to pathological situations such as wounding, fibroblasts become activated and produce ECM components such as collagen I and fibronectin. *In vitro*, coculture of endothelial cells with fibroblasts allows the formation of a perfusable vascular network through self-assembly. Although many *in vitro* vasculature models show that fibroblasts improve vasculature growth, the precise cellular interactions and transcriptional alterations associated with these two interacting cell types are poorly understood. In fact, most data on paracrine interactions between these cell types are gathered from conventional *in vitro* dish cultures, in which both cell types are directly in contact with each other^3–6^. This confounds the results between juxtacrine (direct contact) and paracrine (non-direct contact) signaling. As such, transcriptional analysis of the cells cultured in a precisely controlled microenvironment while being spatially arranged in a non-contact manner will provide important insight into the paracrine cues of fibroblasts affecting vasculogenesis (Fig. 1).

**Fig. 1.**
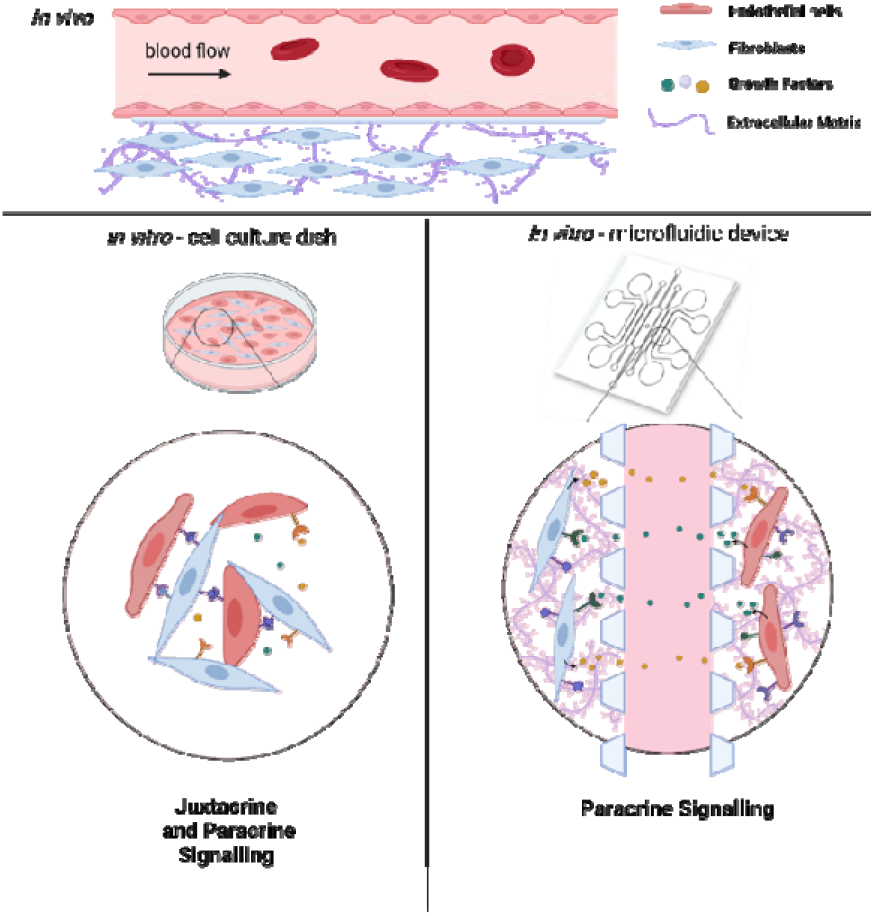
Schematic diagrams showing the *in vivo* condition and the *in vitro* platforms employed to recreate the physiological environment. (Top panel) The *in vivo* environment of blood vessels. (Bottom right panel) Conventional *in vitro* cell cultures such as using a petri dish for 2D culture have both juxtacrine and paracrine signaling which often confounds the results between contact and non-contact interaction.(Bottom left panel) Microfluidic device can be employed to create the partition for these two cell types by separating them in two defined culture area to more accurately and precisely mimicking the paracrine signaling. (Image was created with BioRender.com.)

Here, we employ microfluidics to investigate the paracrine communication of endothelial cells and fibroblasts coculture via a defined non-contact culture arrangement. In this setting, these two cell types are embedded within their respective gel channels and are divided by a media channel, allowing them to only communicate through diffusion. We demonstrate the formation of perfusable and stable vascular networks with accessible open lumens using this approach. The number of cells cultured in this 3D format in the microfluidic device is relatively low, and are embedded in ECM within the microfluidic channels. To analyze the transcriptomic changes of both cell types during the coculturing process, we developed a novel versatile workflow to retrieve cells from the device and extract their RNA^7^. This workflow is widely applicable and useful for downstream analysis of various on-chip 3D cell cultures. Additionally, our transcriptomic analysis with temporal sampling of on-chip interacting cells reveals proangiogenic cues induced by the cocultured fibroblasts, such as growth factors and cytokines, which aligns with our observation that the fibroblasts avoid vessel regression and stabilize vessel formation. We further uncover paracrine signaling molecules that improve vasculature growth through paired receptor-ligand analysis. These paracrine cues could be potentially added into vascular network models where the coculture of fibroblasts in non-contact manner is not desirable. Overall, these findings provide new insights into building more physiologically realistic and precisely controlled *in vitro* vascular models for the integration with organoids as well as various translational applications such as diseases modeling and drug screening.

## Results and discussion

### Fibroblasts prevent vasculature regression and enhance open lumens formation

We employ a microfluidic co-culture platform where the endothelial cells (ECs) are cultured alongside the fibroblasts (FBs) in a separated microfluidic channel, resulting in the formation of a 3D perfusable and functional vascular network^8^. This spatial arrangement allows these two different cell types to interact in a non-contact manner, enabling us to model and probe the paracrine signaling interactions between them. Building on previously reported microfluidic devices^8^, we modified the design to feature seven channels with the central channel potentially used for vascularizing tissue construct (Fig. 2A). The central channel is flanked by two side channels in which the ECs were seeded for self-assembling into the vascular network. The outermost channels of both sides are dedicated for culturing FBs so that any soluble factors can diffuse through the medium channels flanking in between the EC and FB cultures (Fig. S1).

**Fig. 2.**
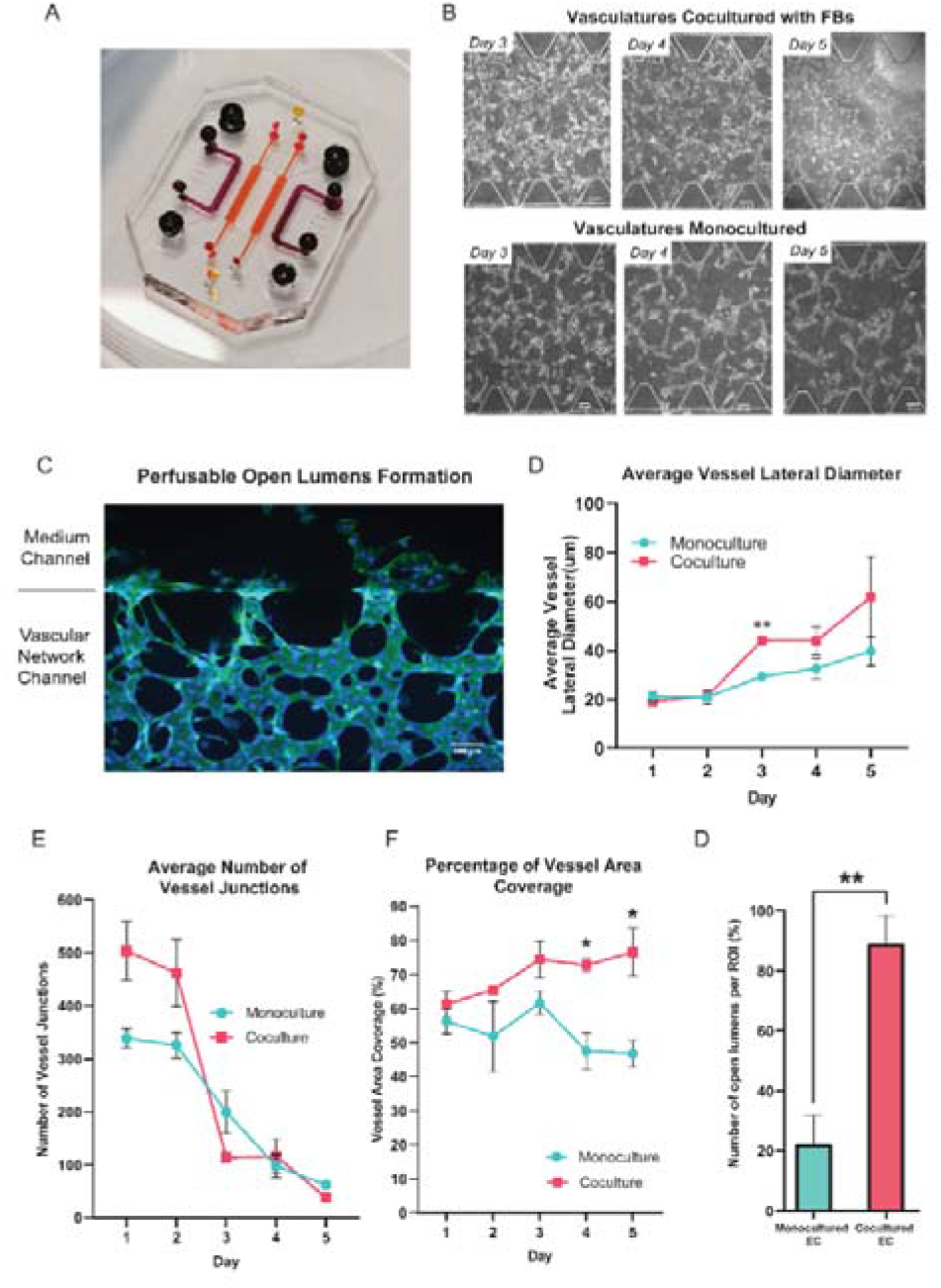
The presence of FBs in a non-contact manner prevents vascular network regression. A)Photograph of the microfluidic chip used in the study to generate perfusable vascular network. Channels are injected with food dye alternately for better visualization. B) Representative images show the comparison of cocultured ECs to monocultured ECs over 3 days. Scale bar =100μm. C) Representative fluorescent image shows the formation of open lumens in which the ECs invade to the other side (medium channel). D) Quantification of the average vessel lateral diameter for both cocultured and monocultured ECs. E) Quantification of the average number of vessel junctions for both cocultured and monocultured ECs. F) Percentage of vessel area coverage over days for both cocultured and monocultured ECs. G)Comparison of the number of open lumens formed between monocultured and cocultured ECs at day 3. *P < 0.05, **P < 0.01, ***P < 0.001, and ****P < 0.0001 (pairwise). (n=3 samples per condition in all cases).

To understand how the presence of FBs affect the vascular morphogenesis in our microfluidic device, we compared the ECs cocultured with FBs to ECs cultured alone (monoculture). In the monoculture condition, either ECs only or FBs are seeded into the device, while in the coculture condition, ECs and FBs are seeded into the device at the same time. During vascular networking formation, single ECs will first elongate in cell morphology, and eventually connect with neighbouring ECs to self-assemble into small vessel buds. Over days, these small vessel buds mature by becoming thicker in diameter, and connecting with other vessel buds to form an assembled network (Fig. S2A). In general, we found the vascular morphology of the cocultured vessel network to be significantly different from the monocultured vessel networks, in terms of the measured average vessels lateral diameter, the average number of vessel junctions, the percentage of vessel area coverage, and the formation and number of open lumens. First of all, both monocultured and cocultured ECs formed branching networks, but the monocultured EC vessels regressed after 4 days in culture (Fig. 2B); In comparison, the ECs cocultured with FBs self-constructed into a vascular network that survived past 4 days with the formation of open lumens that allow beads to perfuse through it (Fig. 2C, and Fig S2B, S2C). During vessel regression, already assembled networks of vessels show ECs with signs of apoptosis such as irregular cell morphology and cell blebbing, and reduced network interconnectedness (Fig. S2D).

Quantitatively, this means that from day 1 to day 3, the number of vessel junctions decreases while the vessel diameter increases for both monocultures and cocultures as the overall interconnectedness of the network improves (Fig. 2D, 2E, S2D). But under monoculture conditions, after day 4 the decrease in vessel junctions reflects increased cell death during network regression rather than improved network assembly (Fig. 2E). The overall condition of the vascular network is best reflected by the vessel area coverage, where a higher coverage area reflects a more robust vascular network. The vessel area coverage continues to increase for the coculture condition, but dramatically decreases for the monoculture after day 3 (From 61.58±3.51μm to 46.75±3.92μm; Fig. 2F).

It is interesting to note that for the monocultured ECs, although vessel regression starts on day 4, the average lateral vessel diameter continued to grow until day 5. This suggests that the ECs in the surviving vessels are not senescent as they are still growing and remodeling, but eventually cannot be sustained. In the coculture condition, however, the average vessel diameter continued to increase all the way from day 2 to day 5 (From 21.31±1.92μm to 61.79±16.35μm). Remarkably, the number of open lumens formed in cocultures is significantly higher than those in monocultures at day 3 even though vascular networks are formed in both conditions at that timepoint (Fig. 2G).

All in all, compared to the EC monoculture, the coculture condition yields larger vessel diameters and a greater number of vessel junctions (see Methods for quantification approach) over a 5-day culture period, and generates a generally healthier and more robust vasculature network with open lumens. Importantly, these results show that fibroblasts can help to prevent early network regression, prolong the lifespan of the vascular network and enhance open lumens formation in a non-contact manner.

### Transcriptomic profiling shows distinct expression profile for monocultured versus cocultured cells at three time points

Having observed and characterized the morphological and functional changes that emerge during vascular morphogenesis of both monocultured and cocultured ECs, we sought to decipher the underlying biological mechanism responsible for the observed phenomenon. We hypothesized that the temporal morphological changes and the phenotypic transitions of the ECs could be governed at the level of transcriptional regulation. In previous studies^9,10^, the RNA expression levels were measured for ECs co-cultured with FBs and ECs monocultured. However, the cells were not separated prior to the analysis, making it difficult to independently analyze the expression changes in individual cell types. Our analysis independently tracks the changes in ECs and FBs for each time point.

To understand at the gene expression level how FBs potentiate and enhance the observed tube-forming ability of ECs in coculture condition, we performed temporal sampling of the ECs and FBs at three time points over the course of cell culturing (Days 1, 3 and 5). Here, we developed a workflow to retrieve the limited numbers of ECs and FBs encapsulated in fibrin gel within the closed channel of the microfluidic device (Fig. 3A; see Methods). After the cells were retrieved, we extracted and purified their RNA, then prepared and sequenced the cDNA library to obtain the transcriptomic profile of ECs and FBs separately for each time point.

**Fig. 3.**
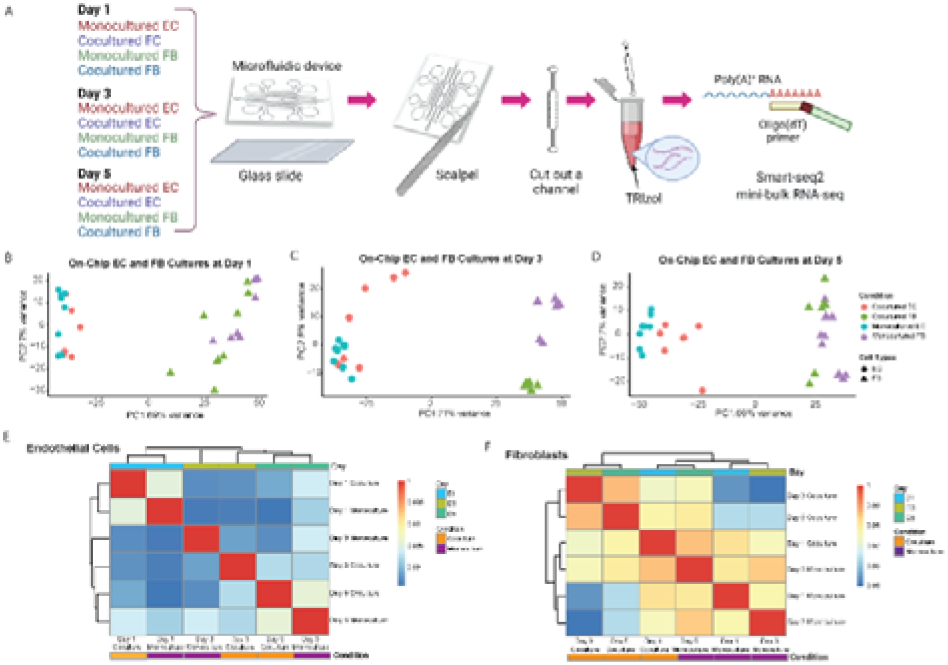
Monocultured and cocultured cells retrieved from microfluidic device show distinct transcriptomic profiles. A) Schematic diagram shows the workflow to retrieve the limited numbers of ECs and FBs encapsulated in fibrin gel within the closed channel of microfluidic device. B) Principal component analysis (PCA) shows the data generated from both monocultured and cocultured ECs and FBs at day 1. The percentage variation in the data explained by the respective principal components is given. Each dot corresponds to individual sample collected from a microfluidic channel. The same applied to Fig. 3C and D. C) PCA plot visualize PC1 and 2 of both monocultured and cocultured ECs and FBs at day 3. D) PCA plot visualize PC1 and 2 of both monocultured and cocultured ECs and FBs at day 5. Cell type confers the greatest degree of variance as shown by the first PC, followed by culture condition effect. E. Correlation heatmap depicting an overview over similarities and dissimilarities between samples. Unsupervised hierarchical clustering groups samples with a more similar gene expression. Samples of endothelial cells can be separated by day, as shown by the annotation bar (Day). F) Correlation heatmap of fibroblast samples showing the separation of monocultured and cocultured fibroblast, as indicated by the annotation bar (Condition).

First, principal components analysis (PCA) on the gene expression data from all the samples across all time points shows that the primary distinction between all samples is the cell type, as PC1 strongly separates ECs and FBs (Fig. 3B-D). Further analysis of the top gene loadings shows genes in PC1 related to EC and FB identity (Fig. S3A-B). This verifies that our cell retrieval workflow recovers transcriptomes of different cell types cultured within a microfluidic confined microenvironment without causing obvious cell type cross-contamination and avoids the time-consuming cell dissociation and sorting that is required for transcriptomic analysis of cells from contact or mixed dish coculture systems. Other than separating cells according to their cell types, the first PC also separates the monocultured ECs from the cocultured ECs, and monocultured FBs from the cocultured ECs for all three different time points (Fig. 3B-D), suggesting that the expression profile of ECs changes in the presence of fibroblasts. Also, we note that the second PC appears to separate the FBs monoculture and cocultures at day 3 due to batch effect (Fig. 3C).

To further check the major sources of variation for all EC and FB groups, we examined the degree of similarity across all conditions and time points by clustering the pairwise correlation of gene expression within the ECs and FBs, respectively (Fig. 3E, F). The ECs are clustered by days, indicating that EC cells at the same stage of maturation are more alike (Fig. 3E). Both monocultured and cocultured ECs of day 1 are clustered together and less similar from the rest, based on the dendrogram distance. While the ECs of day 3 and day 5 are generally similar, the monocultured ECs of these two time points are the most dissimilar within this cluster, indicating that the change in gene expression of monocultured ECs is relatively dramatic from day 3 to day 5. This coincides with our observation that vessel regression in the monocultured ECs occurs starting on day 3. The cocultured ECs, on the other hand, have more stable gene expression from day 3 to day 5 and are transcriptionally more similar to each other. By contrast, the FBs differ more by their culture condition rather than by the days of culture (Fig. 3F).

Overall, these results further support the imaging observations that ECs cultured in the presence of FBs start to diverge morphologically compared to monocultured ECs without FBs at day 3, and become dramatically different at day 5. Having established that the transcriptomic profiles of cocultured cells exhibit notable differences compared to those of monocultured cells, we next sought to identify specific genes associated with these differences.

### Differential gene expression analysis revealed transcriptomic changes in ECs cocultured with FBs at day 5

As evidenced by the morphological and expression profiles of ECs, ECs cocultured with FBs exhibit notable differences compared to the monocultured ECs at day 5. In order to further understand how the presence of fibroblasts may be affecting the ECs in a non-contact manner at this particular time point, we compared the transcriptomes of the ECs under these different culture conditions at day 5 to determine the specific genes that differ in expression with and without FBs, and thereby uncover the biological function of the FBs in this context.

At day 5, we found a total of 490 genes to be differentially expressed between the monocultured and cocultured ECs (P adjusted <0.05, log2 fold change >1). Of these 490 differentially expressed genes (DEGs), 293 genes are upregulated while 197 genes are downregulated in expression for the ECs coculture condition (Fig. 4A). Most of the upregulated DEGs (e.g., LGALS3BP, PCOLCE, FGF5, SEMA3C, CD248, VCAN) in the cocultured ECs have previously been reported to promote vasculogenesis and angiogenesis ^11–16^, suggesting that the ECs exhibited transcriptional changes associated with improved vasculature growth due to the presence of fibroblasts. The full gene list is provided in the supplementary materials (Table S1), shedding light on the potential molecular mediators that enhance the vasculature formation.

**Fig. 4.**
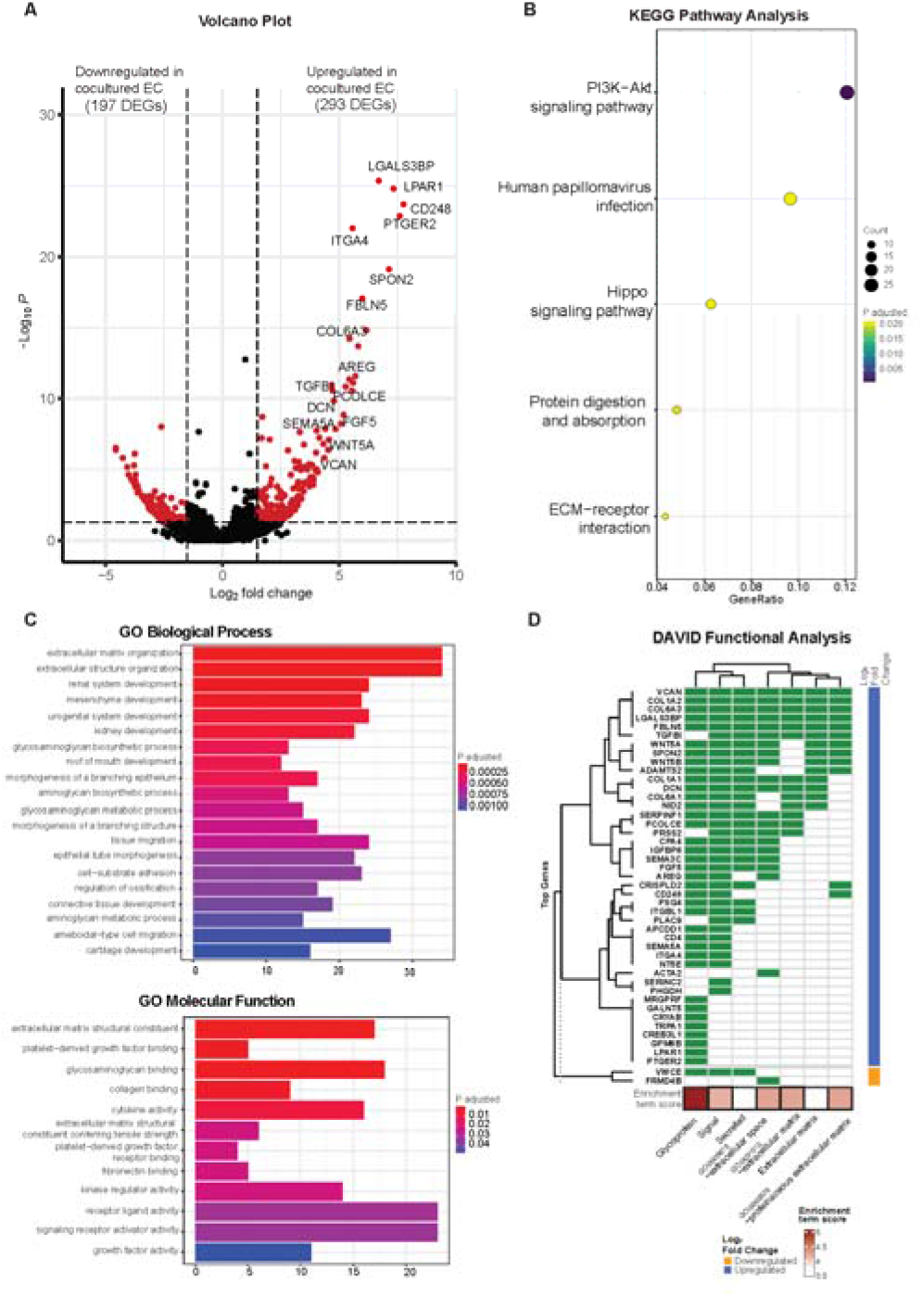
Differentially expressed genes identified for cocultured versus monocultured ECs at day 5. A) Volcano plot shows the DEGs in cocultured ECs versus monocultured ECs. Differential expression is defined by FDR < 0.05 and abs(log_2_FC) >1(red points). 197 DEGs are downregulated in cocultured ECs while 293 DEGs are upregulated in cocultured ECs. B) KEGG pathway analysis displays the significant pathways related to the DEGs identified (FDR < 0.05). C) GO biological processes and molecular functions show the pathways related to both upregulated and downregulated DEGs in cocultured ECs. D) Clustering of DAVID gene enrichment reveals top 75 DEGs are mainly glycoproteins consisted of secreated factors and are located at the extracellular space.

To understand the high-level functions of the identified DEGs, we first performed KEGG and Gene Ontology (GO) pathway analysis, and the results showed that these DEGs are enriched in multiple prominent pathways related to cell proliferation, homeostasis, and cell communication, including the PI3K-Akt signaling pathway, hippo signaling pathway, as well as ECM-receptor interaction, to name a few (Fig. 4B; see Fig. S4A for related genes). Of note, both the PI3K-Akt pathway and hippo signaling pathway are known to promote and regulate cell proliferation and survival as well as angiogenesis ^17^,^18^. GO pathway analysis revealed ECM organization and related proteins binding activity (e.g., collagen, fibronectin), tissues and organs development (e.g., connective tissues, kidney), tissue migration processes, growth factors and cytokines binding activity as well as receptor-ligand activity (Fig. 4C). These are known to be involved in angiogenesis processes. The network plot analysis further showed the DEGs involved in these processes with most of them being upregulated in the cocultured condition (Fig. S4B-C), indicating an overall upregulation in these pathways. This suggests an increase in interactions of ECs with growth factors and cytokines as well as with their surrounding 3D ECM environment when FBs are present, even if the ECs and FBs are non-contact. The large portion of DEGs enriched in various organ development processes suggests links between the functional role of these genes to the organ vascularization process and the potential of using vasculatures formed by coculturing with FB for vascularizing *in vitro* organoid cultures. Many of these DEGs are involved in multiple biological and molecular processes. For instance, procollagen C endopeptidase enhancer 1 (PCOLCE), which has been previously shown to be derived from fibroblast to enhance the lumen formation of vascular networks ^6^, was found to be highly upregulated in ECs coculture and involved in extracellular matrix constituent, glycosaminoglycan binding as well as the collagen binding. This indicates that vessel maturation and stabilization involve a complex set of orchestrated pathways, with proangiogenic cues being one of many key pathways.

Since the GO analysis revealed interactions related to growth factors and cytokines, we also performed the DAVID gene enrichment analysis to check the subcellular locations of the protein products of these DEGs. Remarkably, the majority of the top upregulated DEGs in cocultured ECs are genes encoding secretory proteins and are enriched in cellular components of extracellular space and extracellular region (Fig. 4D), providing additional evidence that coculturing affects the production of secretory proteins that are involved in cell-cell communication.

Our findings identified specific genes and pathways that affect the ECs cocultured with FBs, allowing them to form a more stable vascular network with a longer lifespan than the monocultured ECs, as suggested by prior evidence. Since the top upregulated DEGs in ECs coculture mainly encode secreted factors, we sought to also uncover their respective binding partners in FBs.

### Receptor-ligand analysis reveals paracrine interactions between EC and FB

Receptor-ligand analysis can be used to extrapolate intercellular communication from the coordinated expression of the cognate genes in two heterotypic cells (9). We performed receptor-ligand analysis to further explore how the transcriptomic alterations in cocultured ECs are affected by FBs ^19–21^. Importantly, with the uniquely defined spatial positions of ECs and FBs in the microfluidic device, we could specifically infer and isolate non-contact paracrine intercellular communications and interactions, which is traditionally difficult to tease out from other contact-type interactions ^3,5,9^. Here we specifically looked at the interactions between the ECs and FBs on day 5, since there is the most dramatic difference between the mono- and co-cultured ECs on this day.

There were many receptors or ligands encoded by the DEGs of mono- vs cocultured FBs, whose respective binding partners are also found to be DEGs of mono- vs cocultured ECs. These putative receptor-ligand pairs (RLs) that are differentially expressed in coculture implicate their specific role in the paracrine interactions involved in stabilization and maturation of vascular networks. The receptor-ligand analysis revealed three major classes of signaling molecules that exhibit major differences in expression between mono- and cocultured cells at day 5: Growth factors, extracellular matrix (ECM)-associated factors, and cytokines. Each class of RLs contributes differently to the maturation process.

Growth factors that promote vascular network formation have been widely studied. Among the most commonly discussed and well-validated proangiogenic growth factors are fibroblast growth factor (FGF), platelet-derived growth factor (PDGF), transforming growth factor (TGF-B), and intracellular adhesion molecule 2 (ICAM2), all of which can be secreted by FBs for vascular growth and maintenance ^22^. Our RL analysis also shows interaction of RLs within these families, including FGFR1-FGF5, FGFR1-FGF7, TGFB2-ENG, TGFB1-DCN, PDGFRA-FGFR1, PDGFRB-SIGLEC9, and CD44-ICAM2, suggesting that these pairs have long-range interactions and are not limited to direct contact interaction (Fig. 5A). The growth factors identified in this analysis are consistent with previous studies ^13,22,23^, validating the approach for identifying putative RL pairs between two different cell types. Some growth factors secreted by fibroblasts that we identified in this analysis were not identified before as factors important for vasculature maturation: AREG, B2M, SNX2, SRGN and IGFBP3. These are potential novel factors to be explored as culture media additives or can be functionally added to ECM to support ECs when no FBs are present. We note that cocultured ECs also secreted ligands which not only support their own growth through autocrine signaling but also affect the FBs. The concerted upregulation of FGF5 and FGF7 in ECs and their corresponding receptor, FGFR1 in FBs illustrates the interplay of paracrine cues between both cell types affecting each other. Further mechanistic studies can also be extended to investigate the difference between direct vs non-contact manner of the identified growth factors and their influence on vascular growth and maturation.

**Fig. 5.**
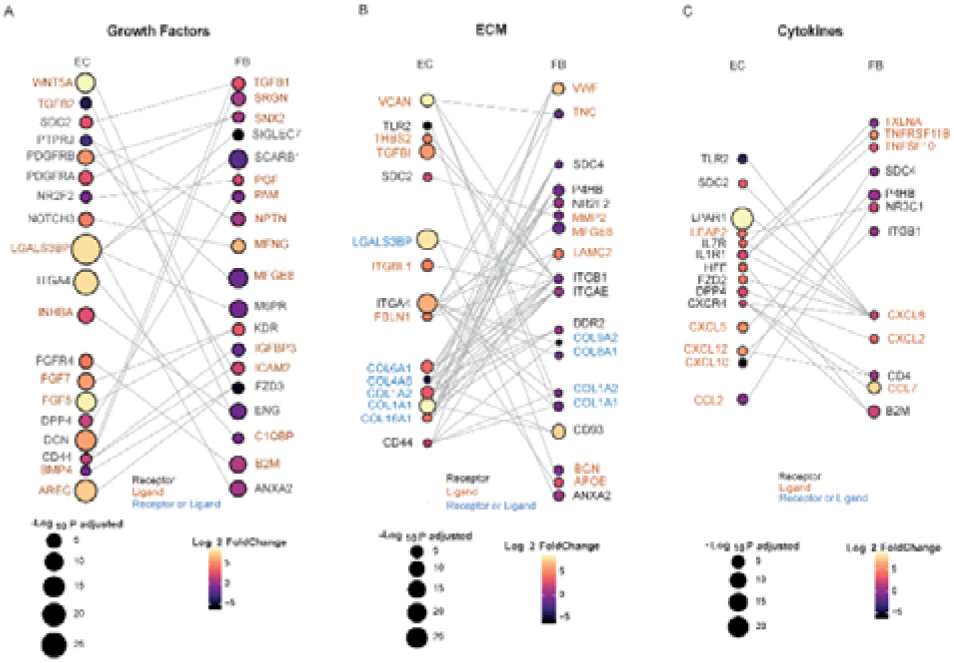
Receptor-ligand analysis reveals paracrine interactions between EC and FB at day 5. A-C. Sankey plots depict the paracrine interaction between the DEGs of EC and FB encoding either putative receptors or ligands, which are grouped into three secretome groups: (A) growth factor; (B) ECM; and (C) cytokines. Color indicates the average log2 fold change of cocultured/monocultured cells, while the dot size represents degree of statistical significance.

One of the major components in the 3D culture microenvironment is the ECM, which can modulate diverse cellular behaviours and structure by constantly undergoing remodeling, whereby the ECM proteins are degraded or deposited ^24^. In addition, ECM remodeling allows branching morphogenesis to occur, which is critical to the formation of vascular networks. The ECM can be modified without intercellular crosstalk, which could explain why the influence of ECs on FBs and vice versa are quite balanced in our analysis for this class of molecules, with ligands and receptors being expressed equally by both cell types (Fig 5B). We found a coordinated expression of ECM related genes by both EC and FB, including ECM ligands such as collagen (COL6A1, COL8A1), fibronectin (FBLN1), tenascin (TNC), proteoglycan (VCAN), matrixins (MMP2) and cell surface receptors such as integrin (ITGA4), tyrosine kinase (DDR2). FBs are known to secrete ECM proteins such as collagen and laminin^25^. Interestingly, majority of putative receptor or ligands related to ECM are upregulated in the ECs coculture condition while downregulated in the FBs coculture condition. This could indicate that the paracrine cue from fibroblasts leads to a general increase in expression of ECM related genes. Contrarily, the ECs could also secrete soluble factors that influence the FBs to down-regulate ECM-associated expression. Notably, most of the collagen genes (e.g., COL1A1, COL1A2) are upregulated in EC coculture yet downregulated in FB coculture. These findings are consistent with previous study that reported the gradual downregulation of collagen synthesis from 3D FB cultures, though the mechanism related to this observation is not clear^26^. Further studies such as proteomics analysis can be performed to understand and compare the deposition of the ECM proteins by the monocultured and cocultured cells^27^.

It is unsurprising that we found many GF and ECM receptor pairs in our analysis, confirming their importance in maintaining a mature vascular network in-vitro. One major new finding of our analysis reveals that cytokines are also a key class of secreted factors in the vascular maturation in coculture systems. Signalling from FBs to ECs include the family of proangiogenic chemokines, such as CXCL5 and CXCL12 which are upregulated in the ECs coculture, while CXCL10, which is angio-static, was downregulated in the ECs coculture. Similarly, signals are also sent from ECs to FBs via cytokine secretion (Fig. 5C): Cytokines such as CCL7, CXCL2, CXCL8, TNFRSF11B and TNFSF10 are upregulated in the FBs coculture, and their corresponding receptors such as CXCR4 are shown to be upregulated in the ECs coculture ^28^. This concerted upregulation of cytokines related receptor-ligands in both cell types further show the significance of their interaction, and this indicates that regulation of inflammatory processes could be very important for regulating the network formation and preventing network regression ^29^. Further studies such as cytokine arrays could be used to verify the relevant soluble proteins and identify their exact roles in promoting vascular growth.

Overall, we identified concomitant receptor-ligand pairs that are involved in the paracrine communication between ECs and FBs. Although further functional validation of their putative interaction, these results can have a considerable impact on tissue engineering of more realistic vasculatures *in-vitro*.

### Temporal transcriptional changes between ECs monocultured vs ECs cocultured with FBs

While we have identified the putative receptor-ligands of both spatially controlled cells, we were also interested in investigating their temporal gene expression patterns to study if the gene expression of the specific receptor or ligands change in a time dependent manner. We performed likelihood ratio test to look at the transcriptional changes of these genes across different time points. Remarkably, our analysis shows that 27 EC genes and 12 FB genes encoding receptors or ligands display statistically significant changes across days (Fig. 6 and Fig. S6-S7), corroborating our findings of these genes as putative receptors and ligands responsible for improved vasculature growth. The EC genes identified mostly have higher expression in cocultured condition, including ECM related genes (COL1A1, FBLN), chemokines related genes (CXCL10, CXCL12) and growth factors (FGF5, AREG) (Fig. 6A-F). Notably, COL1A1 has been indicated to play a role in EC lumen formation (Fig. 6A); and CXCL12 has been recently found to enhance angiogenesis in a dose-dependent manner^30^ (Fig. 6E). These ligands could be potentially supplemented into the medium of vasculatures to replace fibroblast culture. Interestingly, majority of the statistically significant genes of FB cultures have lower expression across days for the cocultured condition (Fig. 6G-I), including TNC that has an inhibitory impact on vascular sprouting and contrastingly induce pro-angiogenic factors in the presence of cancer cells^31^,^32^. Several of these genes, ENG, BGN and IGFBP3 that have been implicated to promote angiogenesis^33–37^ are found to have significantly lower expression temporally, suggesting that they might be depleting in FB gradually over time. It is likely that these genes promote vasculature growth at the beginning, and the further addition of these identified ligands into the culture system at the later stage of vasculature cultures could potentially enhance the vasculature growth and prolong their lifespan. Together, these temporal studies validated the paracrine receptor and ligands identified and indicated their potential requirements in a time-dependent manner.

**Fig. 6.**
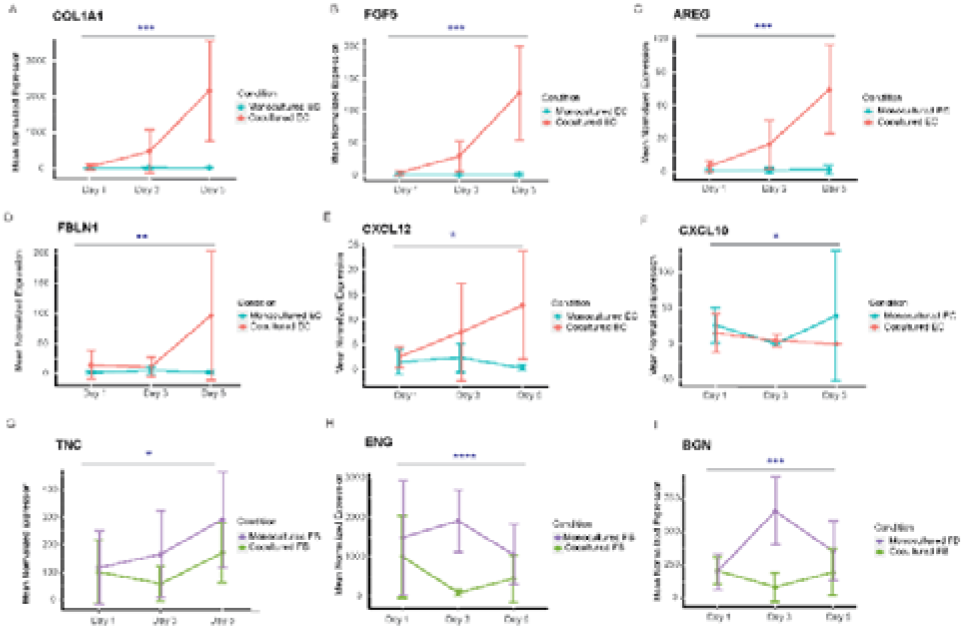
Temporal gene expression patterns of genes encoding receptor or ligands. A-F) Representative line plots showing the trend of DEGs (P-adjusted <0.05 in temporal analysis) encoding ligands of ECs in both cocultured and monocultured condition. G-I) Representative line plots showing the trend of DEGs (P-adjusted <0.05 in temporal analysis) encoding ligands of FBs in both cocultured and monocultured condition. Likelihood ratio test are performed for all the graphs. *P < 0.05, **P < 0.01, ***P < 0.001, and ****P < 0.0001.

In addition, we also look at the dynamic trend of a group of genes that show transcriptomic changes for cocultured versus monocultured cells across three time points. Hierarchical heatmap show the differences in expression between cocultured and monocultured ECs across three days, which were further categorized into four different clusters (Fig. 7A), in which cluster 1 are mainly genes that have higher expression levels in EC coculture condition at day 5, and are associated with pathways related to the extracellular matrix, cell-substrate adhesion and tissue developments (Fig. 7B) while cluster 3 are genes that start to be highly expressed for EC coculture at day 3 with the majority of them staying highly expressed until day 5. GO biological pathways analysis show that this cluster of genes are largely enriched in pathways similar to cluster 1, including extracellular matrix and organs development (Fig. 7C). Interestingly, the group of genes in cluster 4 with higher expression at both day 1 and day 3 are shown to be associated with cytokines and chemokines related pathways (Fig. S8). This suggests that different pathways govern the changes of cocultured ECs temporally in which ECM remodelling and organ developments largely occur from day 3 to day 5 while the pathways related to cytokines and chemokines started earlier from day 1.

**Fig. 7.**
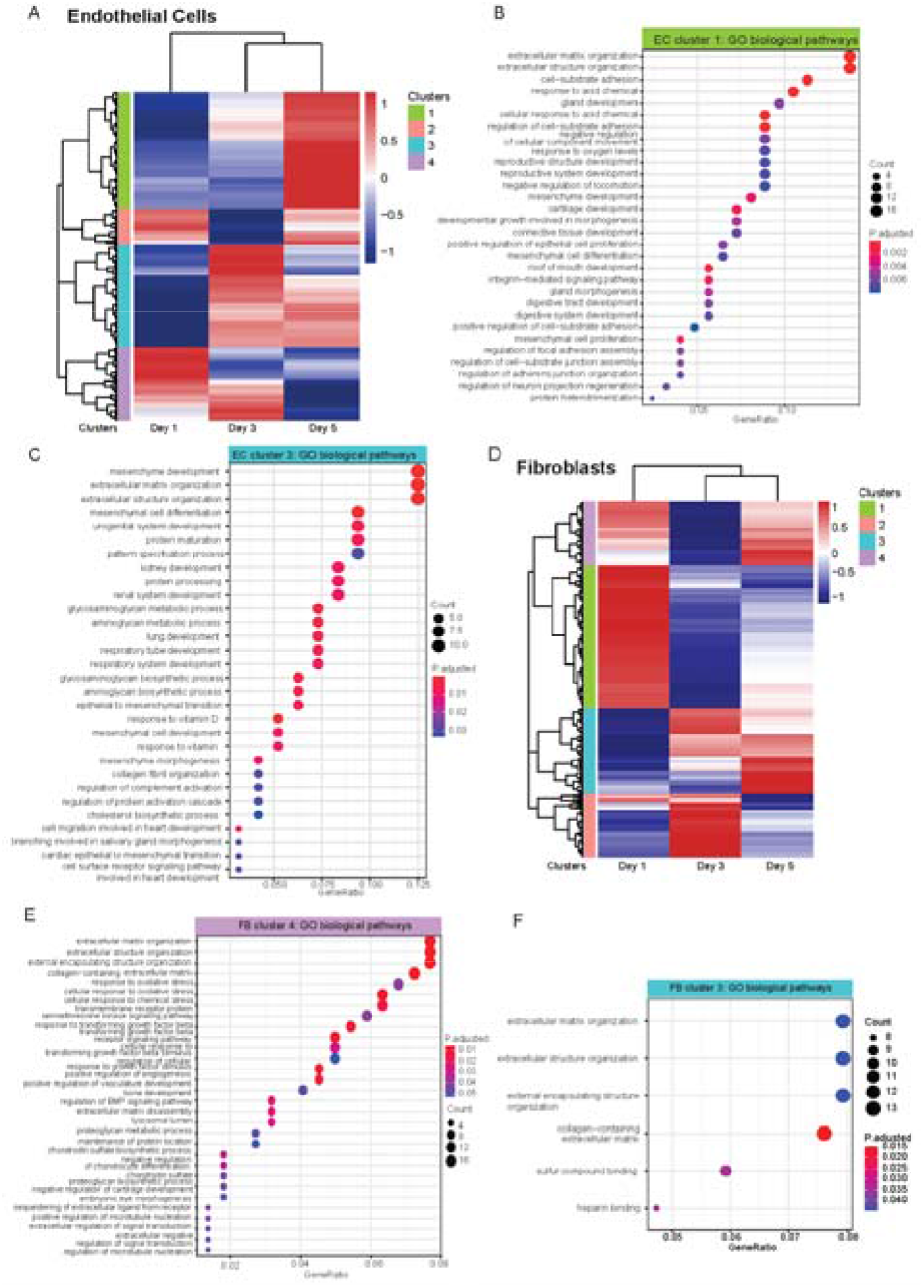
Temporal transcriptional changes between monocultured and cocultured cells. A) Heatmap showing the temporal transcriptional changes between monocultured and cocultured ECs, which can be grouped into 4 different clusters. B) Go biological pathway analysis displays pathways associated with EC cluster 1. C) Go biological pathway analysis displays pathways associated with EC cluster 3. D) Heatmap showing the temporal transcriptional changes between monocultured and cocultured FBs, which are grouped into 3 main clusters. E) Go biological pathway analysis displays pathways associated with FB cluster 4. F) Go biological pathway analysis displays pathways associated with FB cluster 3.

We also performed the same analysis for FB cultures to look at the temporal changes in their gene expression between cocultured and monocultured FBs, visualizing the results in a heatmap with four clusters. Cluster 4 are genes enriched in pathways associated with ECM, vasculature development as well as responses to environmental stimulus and growth factors, and they generally have higher expression in day 1 and day 5 (Figure 7E), suggesting that FBs supported the vascular network formation temporally through the ECM remodelling and growth factor signalling. Similarly, cluster 3 consists of genes significantly enriched in ECM organization, and they are genes with lower expression in cocultured FBs at day 1 while increasing in expression at day 3 and day 5, indicating that ECM deposition, remodelling or degradation might largely occur on these two days (Fig. 7F).

These findings suggested that ECs and FBs not only interacted in a paracrine manner through a spatially controlled location but also temporally with different transcriptomic changes across days when compared to monocultured conditions. These also correlates with our initial results that showed that FBs caused critical phenotypic changes to ECs as compared to monocultured EC at day 3, and that FBs avoided network regression that greatly segregate both conditions at day 5 wherein an interconnected perfusable network formed in coculture and random sprouts with collapsed lumens formed in monoculture condition.

## Experimental Methods

### Microfluidic device design and fabrication

The microfluidic devices used in this study were made of polydimethylsiloxane (PDMS) and modified from a previous study^8^. They are comprised of seven parallel channels with dimensions outlined in detail (Fig. S1). Soft lithography and replica molding were used to fabricate the devices. Briefly, a lithography mask was prepared based on the AutoCAD design. A silicon mold was fabricated out of SU8-2050 photoresists (Microchem) using the standard photolithography techniques. PDMS (Sylgard 184 Silicone Elastomer, Dow Corning) was mixed at a 10:1 ratio of base to curing agent to be poured over the silicon master. Then, the PDMS was allowed to cure for at least 2 hours at 80°C before being removed from the master. The hydrogel injection ports and the reservoirs for the cell culture were punched using the PDMS punchers. Trimmed PDMS devices and glass coverslips (BBAD02400400#A1, 24mm x 40mm, Thermo Fisher Scientific) were then separately cleaned with IPA, distilled water, scotch tape, and air gun before being treated with oxygen plasma for 3 minutes to form an irreversible bonding. The devices were then incubated overnight on hotplate at 80°C and sterilized by UV irradiation for at least 30 minutes before each experiment.

### Cell culture

Human umbilical vein endothelial cells (ECs, CC-2519, Lonza) were cultured in Endothelial Microvasculature Growth Medium (EGM-MV BulletKit^™^, CC-3202, Lonza), and normal human lung fibroblasts (NHLFs, CC-2512, Lonza) were grown in Fibroblast Growth Medium (FGM-2 BulletKit^™^, CC-3132, Lonza). Cells were cultured under a humidified incubator at 37°C and 5% CO^2^ and grown up to 80% confluency for experiments. All the cells no later than passage 8 were used for the experiments.

### Gel preparation and cell seeding

Gel preparation was performed as previously described^8^. To generate the fibrin gel, the fibrinogen solution and thrombin solution were prepared first separately. Briefly, the fibrinogen solution was prepared by dissolving 15mg of bovine fibrinogen (F8630-1G, Sigma-Aldrich) in 2.5mL of PBS in a 37°C water bath for at least 1 hour, which was then further sterile filtered using a 0.22μm filter. Diluted thrombin solution was prepared by adding thrombin stock solution (T9549, Sigma-Aldrich) to culture medium to have a final concentration of 4U/mL. Each channel of the microfluidic devices was also rinsed with DPBS before cell seeding. For cell seeding, ECs and FBs were trypsinized using 0.25% Trypsin-EDTA (25200056, Thermo Fisher Scientific) for 2 minutes, neutralized with the EGM-2MV medium and then centrifuged for 3 minutes at 200g. Media was then aspirated from the cell pellets, and the cell solutions were then put into the thrombin solution before quickly mixing with 6mg/ml fibrinogen by pipetting up and down for a few times. The cell-laden gel mixtures (6 million cells per ml of ECs and 3 million cells per ml of FBs) were then gently introduced into their respective channels and allowed to polymerize for 15 minutes at room temperature by putting them into a humidified chamber. Once polymerized, the channels of the cell culture medium were then loaded with medium. All the devices were then incubated under a humidified incubator at 37°C and 5% CO_2_. Every 24 hours the cell culture medium was removed and refilled with fresh EGM-2MV medium.

### Immunostaining and imaging

To visualize the vascular network morphology on day 5 in which they are fully formed, standard immunostaining protocol was used to fix the ECs. After aspirating the medium from the medium channel, 3D vasculatures in the device were washed with PBS solution three times. On-chip vasculatures cells were then fixed with 4% paraformaldehyde (PFA) in PBS solution for 15 minutes and permeabilized with 0.1% Triton X-100 in PBS solution for another 15 minutes. They were then blocked with 3% bovine serum albumin (A9418-500G, Sigma Aldrich) in PBS for 2 hours at 4°C. Following washing, Phalloidin (1:200, Alexa Fluor488, Invitrogen) and DAPI (1:1000, Invitrogen) in PBS were added into the devices, which were then incubated overnight at 4°C. Next day before imaging, all samples were washed three times with PBS before imaging. Confocal microscopes (Leica, SP8) and fluorescent microscope (Zeiss) were used for imaging. Images were processed by using Las X imaging software (Bitplane, Belfast) and ImageJ (NIH).

### Quantification of the vascular network

To quantify the vascular network formation and stabilization, phase contrast images were preprocessed and several parameters such as the coverage area of vessels, the number of vessels and the lateral diameter of the vessels were computed using AngioTool (NIH) and Fiji distribution of ImageJ (NIH) as described previously^38–40^. Lateral diameter was computed as projected lateral vessel area divided by total vessel length^38^. Interconnectedness is defined by the number of vessel endpoints divided by the number of vessel junctions, and the value of interconnectedness close to 0 indicates good interconnected vascular network^41^.

### Vascular network perfusion

To confirm that the vascular networks formed in our device are functionally perfusable, fluorescent polystyrene beads of 5 μm in diameter suspended in PBS solution were introduced into the vasculature. All the beads were treated with 3% bovine serum albumin (A9418-500G, Sigma Aldrich) to prevent aggregation of the beads. On day 5 of the vasculature formation and cell culture, the reservoirs of each device were aspirated, and the beads in PBS solution were added on one side to create a hydrodynamic imbalance to induce fluid flow through the vasculature. The vasculature was then observed using microscopy as described in the imaging section.

### RNA isolation of 3D cells and tissues from microfluidic device

Microfluidic devices for this application were prepared with some modifications made in the fabrication process. Reversible bonding techniques was applied, and glass slides were used for the bonding of the devices. Briefly, the device was pressed hard onto the glass slide and then incubated overnight on a 120°C hotplate. The device was then used for culturing as described earlier. To extract the RNA of the cells embedded in fibrin gel, we peeled the reversibly bonded device off the glass slide slowly. The fibrin gels occupied with cells or tissues remained inside the channels of the detached microfluidic device. Since different cell types were being cultured in separated channels, we carefully used a scalpel to cut out the individual channel and rinsed it in TRIzol reagent (15596026, Thermo Fisher Scientific) for 20 minutes. This avoided different cell types from being mixed and contaminated. RNA isolation along with DNase application (M0303, New England Biolabs) were performed to purify the samples.

### Library construction and sequencing

Mini-bulk cDNA library was constructed using Smart-seq2 protocol that uses oligoDT to prime the polyA RNA for cDNA production^42^. The concentrations were quantified by Qubit 3.0 fluorometer (Thermo Fisher Scientific) and cDNA library size was checked using Fragment Analyzer HS NGS Fragment Kit (1-6000bp) (Agilent formerly Advanced Analytical). Only high-quality cDNA libraries without notable degradation were used for sequencing library construction. Illumina sequencing libraries were prepared using Nextera XT DNA Library Prep Kit (Illumina). The concentrations of samples were diluted to 0.1-0.3ng/ul. Tagmentation and dual index adding were done according to the C1 Fluidigm protocol. The library samples were then pooled with equal volumes and sequenced using Nextseq500/550 Middle Output Kit v2.5 on Nextseq500 sequencer (Illumina) to produce paired-end 75 base long reads.

### Data processing and analysis

Quality of all raw FASTQ files was checked using Fastqc, and reads were aligned to the human reference genome GRCh38 using STAR with default parameters^43^. Read group BAM files were merged together using SAMtools^44^, and gene counts were made using featureCounts^45^. PCA and unsupervised hierarchical heatmap clustering were performed using the DESeq2 package^46^. For the hierarchical heatmap clustering, the degree of similarity was checked for each cell type by using the average log transformed normalized gene expression of all biological replicates. Top gene loadings of PCA were further analyzed using PCAtools. Combat-seq was used to remove batch effect among different sample batches when necessary^47^. All differentially expressed genes in the transcriptome data were identified using a generalized linear model with the Wald statistical test in DESeq2, and with the assumption that gene expression count data were dispersed per a negative binomial distribution. The P-values were adjusted using the Benjamini and Hochberg’s approach for controlling the false discovery rate. Volcano plot was performed to infer the overall distribution of DEG using the EnhancedVolcano package. Gene ontology (GO) enrichment analysis and Kyoto Encyclopedia of Genes and Genomes (KEGG) pathway enrichment analyses were performed with the R package clusterProfiler and g:Profiler using p < 0.05 as the threshold^48,49^, and the DAVID gene annotation system was used to check subcellular locations of the genes. Paired receptors-ligands were selected using database from previous publications^19^,^21^ and temporal gene expression patterns were performed using DESeq2 package with likelihood ratio test (LRT). Dot plots and line plots were plotted using ggplot2 package. All analysis were performed in the R Statistical Computing environment v3.3.1.

### Statistical analysis

Time series line plots and bar charts are plotted as mean±SD with the software GraphPad Prism 9.2.0, unless indicated otherwise. RM two-way ANOVA with Bonferroni multiple comparison test was performed for the comparison of two culture conditions over days. Significance is represented as follows: \**P* < 0.05, \*\**P* < 0.01, \*\*\**P* < 0.001, and \*\*\**P* < 0.0001.

## Conclusions

Previous studies have shown the formation of vascular network in the presence of fibroblasts as supporting cells^8^,^50^,^51^. Our findings corroborated their reports that coculturing with fibroblast can avoid vascular network regression and improve vessel lumen formation. Although these studies demonstrated that fibroblasts play an important role in maintaining vessel integrity and survival, the fibroblast-derived soluble factors that play important roles in angiogenesis have not been fully explored yet. Progress in this research area has been hampered by a lack of proper *in vivo* and *in vitro* models of the physiological environment of vasculatures. Previous transcriptomic studies related to vasculature have been largely conducted without a curated microenvironment, which is now provided by the microfluidic platform^12^. Specifically, traditional *in vitro* platforms lack the ability to perfuse the vasculatures and lack proper spatial arrangement of cells, meaning that crosstalk between ECs and FBs can only be assessed through a direct contact manner, which is more like juxtacrine signaling: Juxtacrine signaling involves cells with direct physical contact while paracrine signaling is a form of cell-cell communication in which cells produce signals to stimulate changes in neighbouring non-contact cells. A degree of oversimplification inherent in many of these cultures without the proper patterning of cells leave many important questions unresolvable. The fibroblast cues to control angiogenesis in non-contact manner remain understudied, and the transcriptional response of perfusable vasculature to fibroblast as well as 3D ECM environment has not yet been explored in-depth. Addressing these questions are essential for being able to artificially replicate an optimized vascular culture environment robustly and with control.

We demonstrate here the elucidation of true non-contact paracrine communications between endothelial cells and fibroblasts cocultured via a microfluidic platform. In this setting, these two cell types are embedded within their respective gel channels and are divided by a media channel. Thus, they communicated only through non-juxtacrine diffusion of soluble factors via the medium channel between them, resulting in a more stable and matured vascular network in comparison to the monocultured condition. It is also challenging to retrieve these low numbers of cells from microfluidic device, and it is even harder with cells encapsulated by fibrin gel. Conventional approaches used to study the transcriptome profile of 3D cell culture or tissues from microfluidic device either requires scaling up the microfluidic device to get more cells, or using the 2D state to represent the 3D state of the cells as it is easier to extract the DNA or RNA from 2D cultures. Thus, we also developed a simple method to extract the RNA from our vasculature out of the microfluidic device. We showed that our technique in retrieving the transcriptomes of on-chip cells can overcome the limitations to perform downstream studies of 3D gel encapsulated cells in the microfluidic device.

Through the transcriptomic profiling and comparison of monocultured versus cocultured ECs and FBs, we identified many key growth factors and cytokines that could improve vasculature growth. For example, we identified VCAN, LGALS3BP, CD248, PTGER2, ITGA4, SPON2, SEMA3C and COL1A1, all of which have been implicated in wound healing and extracellular matrix processes. GO and KEGG pathway analyses further implicate these upregulated DEGs in cell proliferation and morphogenesis as well as tissue development related processes, indicating the role of these gene products in vessel maturation, given the study context. Furthermore, closer inspection of the DEGs revealed many genes encoding secretory proteins, thus, we performed putative receptor ligand analysis to examine the non-contact receptor-ligand interactions. We found three groups of paired receptor-ligand analysis: growth factors, ECM-associated factors, and cytokine related receptor-ligand pairs. The related ligands identified such as VCAN, VCAM1, TNC may be useful in replacing fibroblast cultures to support the vasculatures maturation. These paracrine cues were supported by our temporal analysis showing significant changes in their transcriptional levels for monocultured vs cocultured conditions over days. Based on these results, we posit that FBs are needed throughout the 5 days of the vasculature growth: they are involved in the vasculature morphogenesis process starting from day 1; trigger the inflammatory and cytokines related pathway at day 1 and day 3; and stabilize the vessels as well as ECM at day 3 and day 5.

In summary, *in vitro* microfluidic based vasculature model is a unique platform that allows proper spatial arrangement of multiple cell types in the same microenvironment. With this model, we developed an approach to transcriptionally interrogate the paracrine interaction of two interacting heterotypic cells, namely ECs and FBs. We further uncovered the set of proangiogenic cues that can be potentially added into the medium system to improve *in vitro* vascularization models. Putative paired receptor-ligand studies further showed that FBs regulate vasculature formation through multiple interactions. This also suggested a list of proangiogenic factors that could potentially replace FB in the culture system although future studies would be needed to test out different combinations of angiogenic factors required to achieve this. The role of cytokines and the inflammatory process during vessel maturation and remodeling is an important direction of study which remains difficult to perform using 2D cultures. Many have attempted to study the interaction and effect of immune cells such as macrophages on vessel formation, growth, maintenance, and remodeling. But vessel perfusion, which serves as one way for immune cells to gain access to the vessel, is not possible in 2D formats, making it challenging to realistically replicate the *in vivo* interaction of these two cell types in conventional cultures. With a microfluidic platform like the one we use here, coupled with our cell retrieval method and comprehensive transcriptomic analysis, it will be possible to investigate vessel-immune interactions. Our present study serves as a framework for future investigations of complex cellular interactions.

## Author Contribution

Conceptualization: SYT, AW

Methodology: SYT, QYJ, ZL, YX

Data Analysis: SYT, QYJ

Supervision: AW

Writing—original draft: SYT, AW

Writing—review & editing: SYT, AW

## Acknowledgements

For helpful discussion and suggestions, we thank: Prof. Hongkai Wu and Prof. Fei Sun at the Hong Kong University of Science and Technology (HKUST); Prof. Noo Li Jeon at Seoul National University and his lab alumni Dr. Somin Lee. We also thank Sukey Chan and Tom Benavides for the help in device fabrication; Alex Fung for the valuable insights in device troubleshooting; Lily Cheng for the help in cell culture; and support staffs of the Biosciences Central Research Facility and the Nanofabrication Facility at HKUST for the help in sequencing and device fabrication. We also thank all the members of the Wu research group for helpful discussions and administrative assistance during this project.

