## Supplemental Figures and Legends for "Transcriptomic Analysis of 3D Vasculature-On-A-Chip Reveals Paracrine Factors Affecting Vasculature Growth and Maturation"

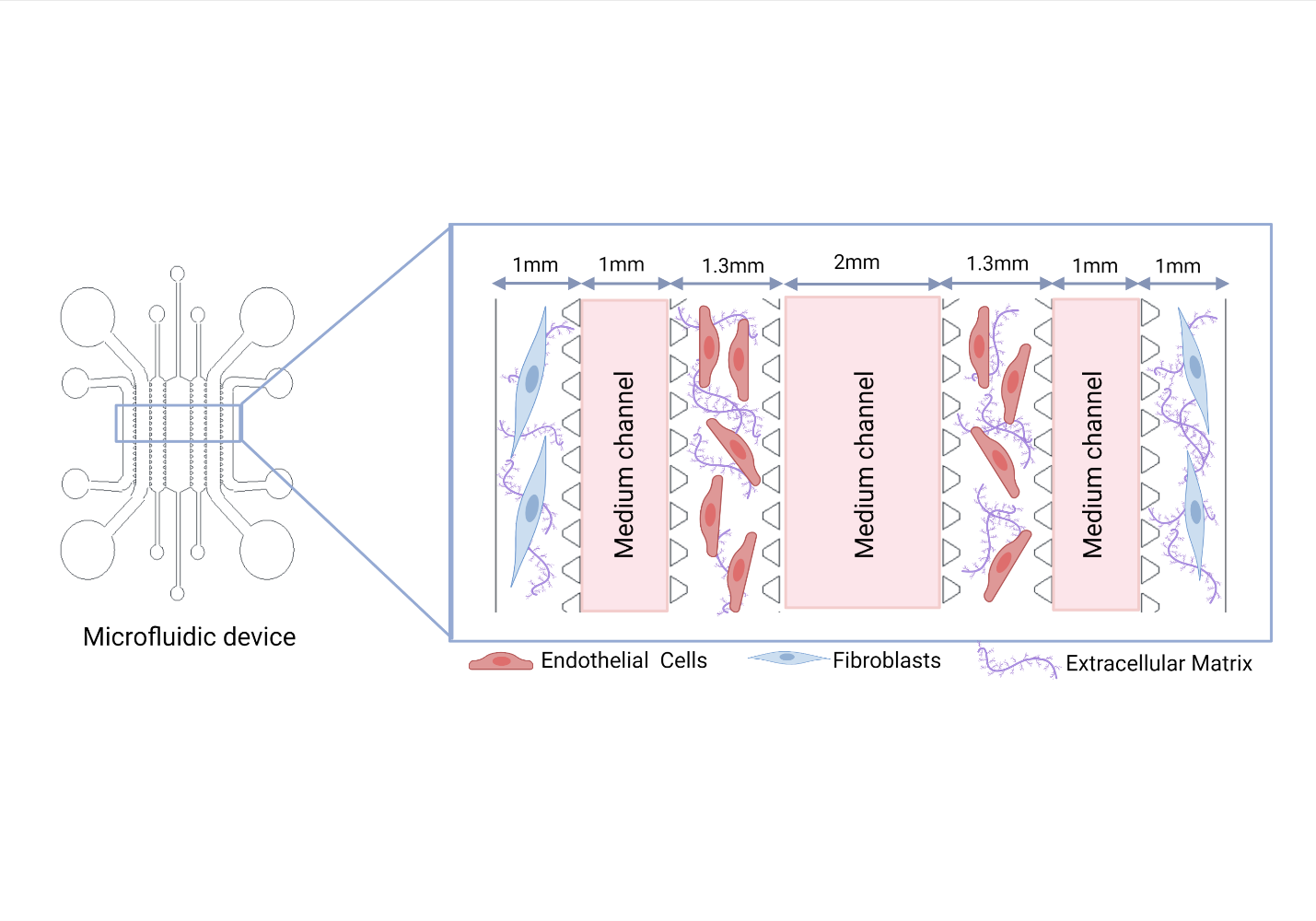

**Fig. S1. Schematic showing the details of the microfluidic device.**

The microfluidic device is designed to have seven channels. All the channels are partitioned by microposts to separate different cultures and prevent the leakage of the gels to other channels while allowing medium to flow through. An enlarged schematic gives a better view of the microposts and the dimensions of the channels. Channel height: 120 - 150μm. Pictures shown here are not scaled. (Created with BioRender.com.)

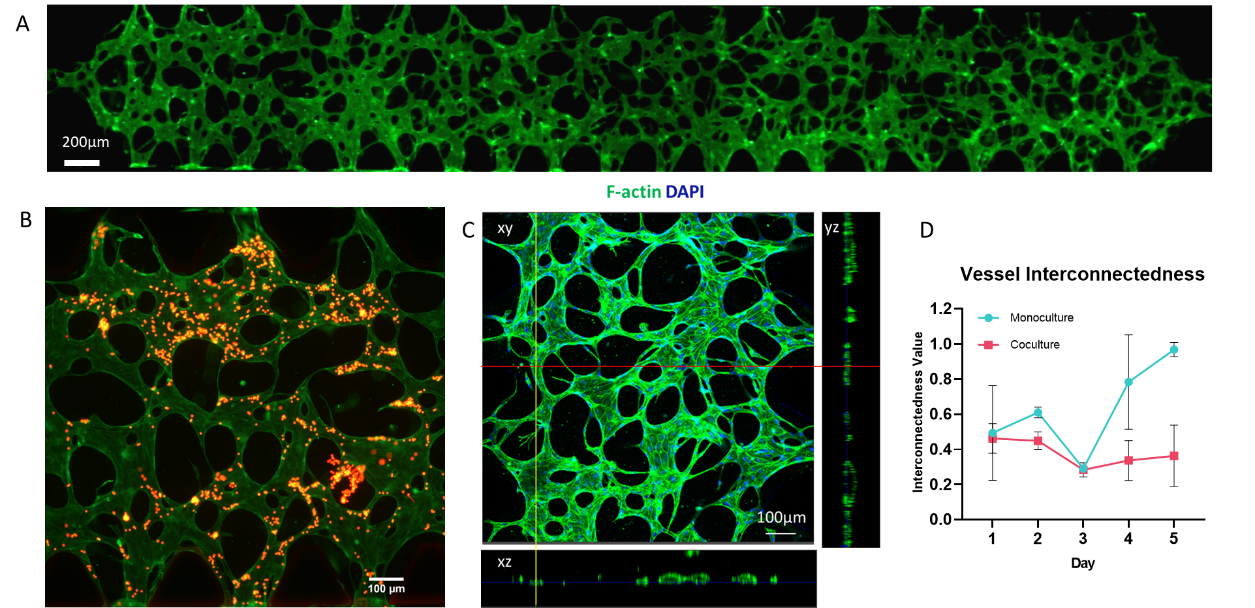

**Fig. S2. Formation of 3D perfusable vascular network by coculturing with fibroblasts**
A. Stitching of images showing the vascular network spanning across the channel, scale bar =200um.

B. Perfusion of 5um fluorescent beads through the vascular network, scale bar = 100um.

C. Fluorescent images displaying the immunostaining of the vasculature formed. 2D fluorescent images projected from 3D z-stack showing the vascular network and hollow lumens formation at day 4; z-stack: 180µm, scale bar = 100µm.

D. Line graph tracking the interconnectedness of vascular network monocultured vs cocultured with FB from day 1 to day 5. Interconnectedness value defined as the number of vessel endpoints divided by the number of vessel junctions.

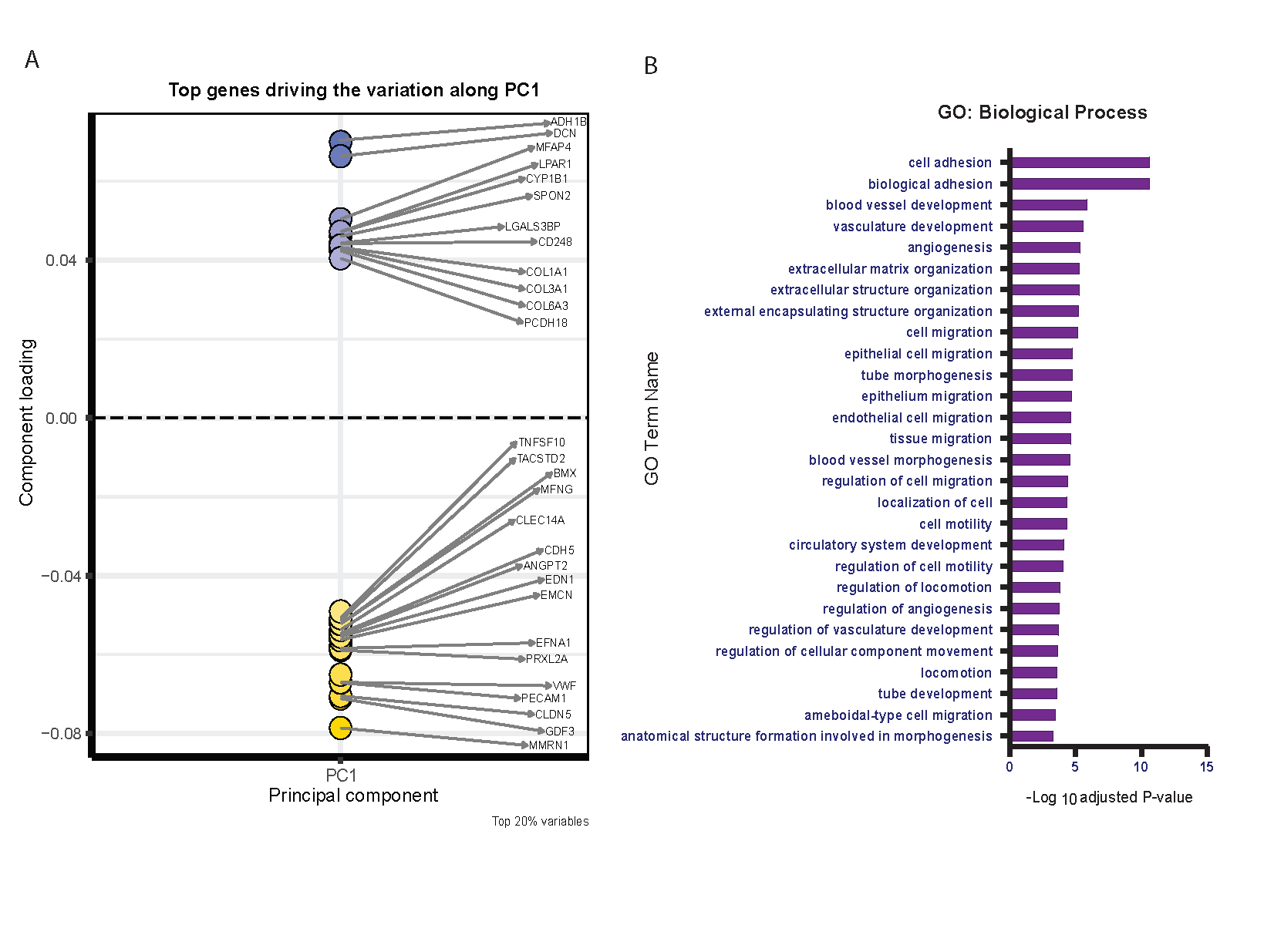

**Fig. S3. Top gene loadings shows genes in PC1 and PC2.**
A. Representative component loading plot displaying the top genes driving the variation along PC1 that separate EC and FB as two different cell types. The blue dots corresponds to genes related to FB identity while yellow dots corresponds to genes related to EC identity.

B. GO biological process analysis shows pathways related to the biological function of EC and FB.

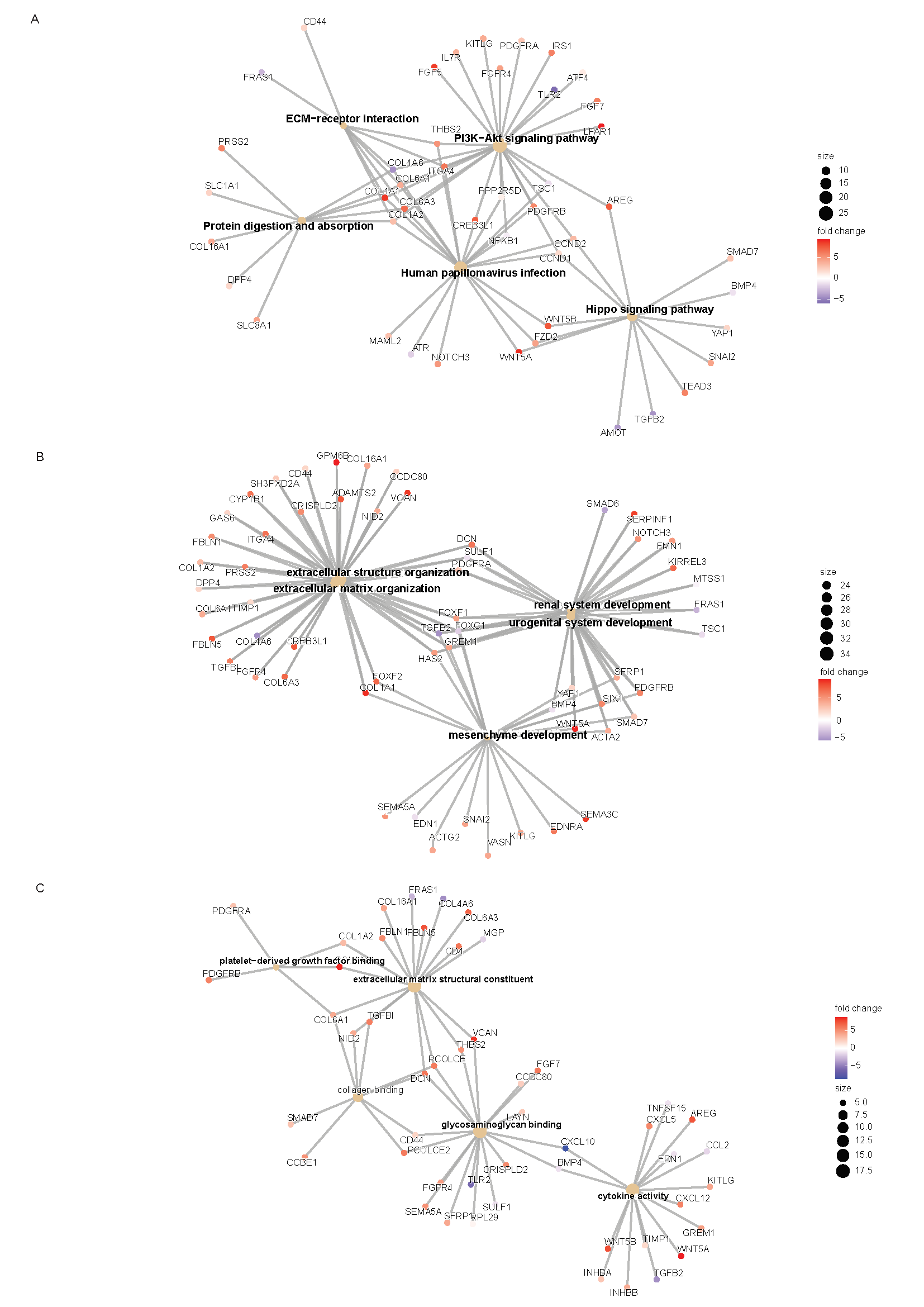

**Fig. S4. Network plot shows the DEGs involved in** A. KEGG pathway; B. GO biological processes and C. GO molecular functions, respectively.

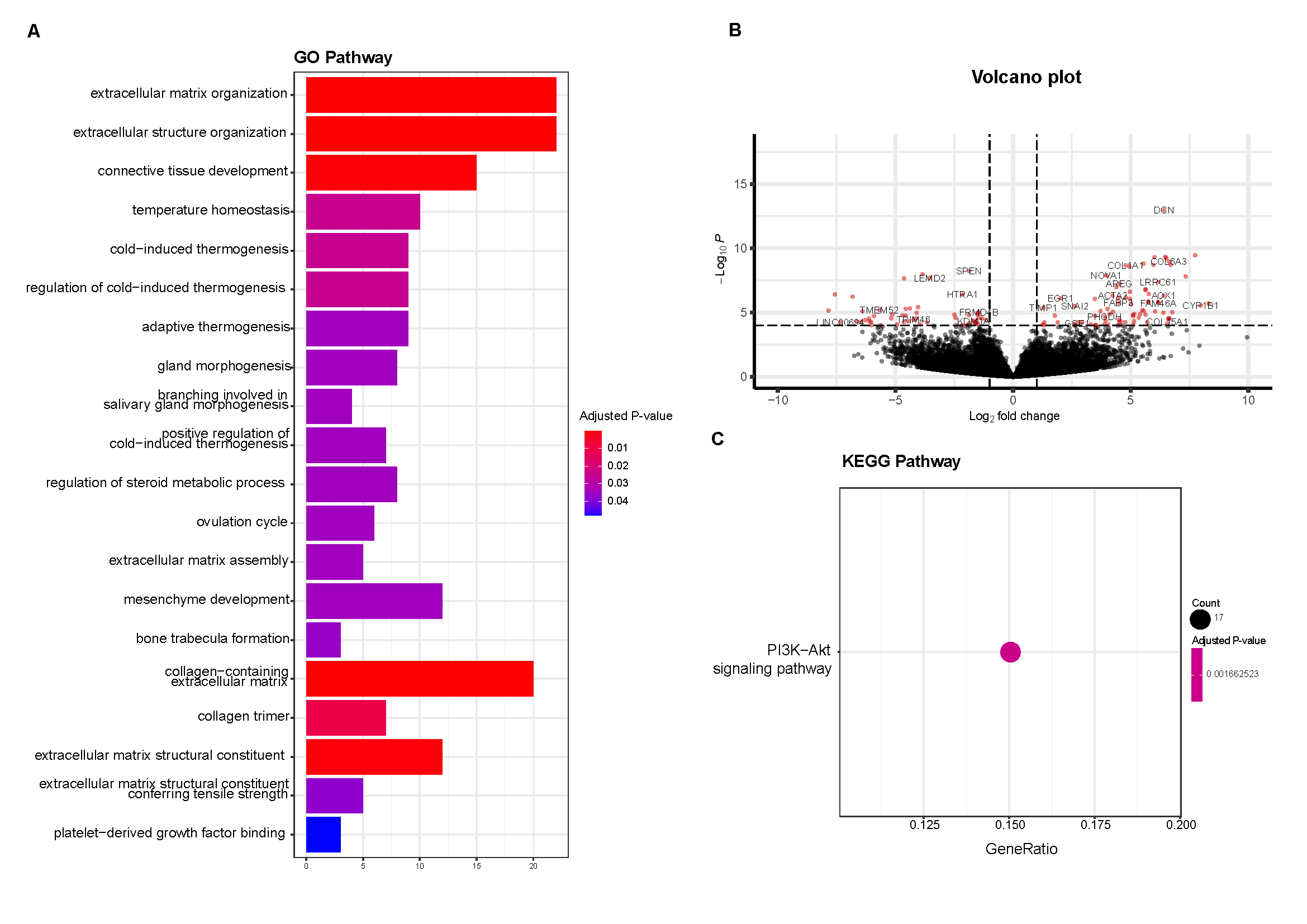

**Fig. S5. Differentially expressed genes identified for cocultured versus monocultured ECs at day 3**
A. GO pathway analysis show the pathways related to both upregulated and downregulated DEGs in cocultured ECs at day 3.
B. Volcano plot shows the DEGs in cocultured ECs versus monocultured ECs at day 3. Differential expression is defined by FDR < 0.05 and abs(log2FC) >1(red points). 
C. KEGG pathway analysis displays the significant pathways related to the DEGs identified (FDR < 0.05).

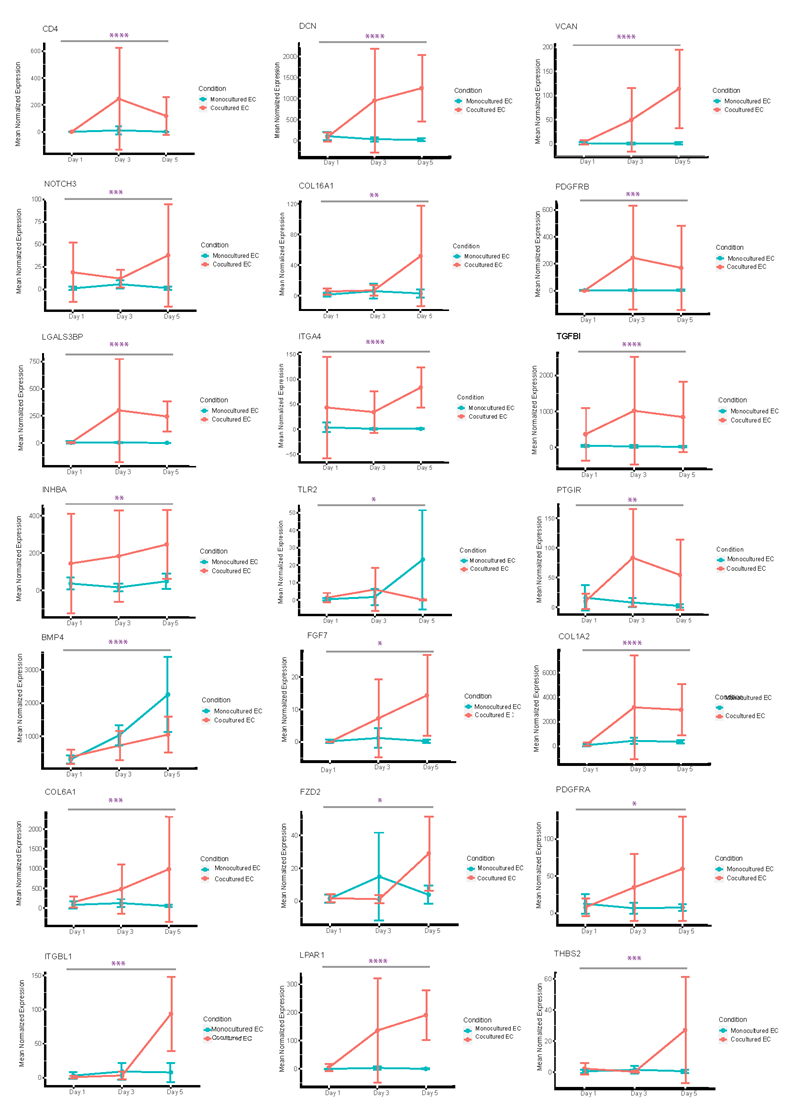

**Fig. S6. Temporal gene expression patterns of EC genes encoding receptor or ligands.**
Line plots showing the trend of DEGs (P-adjusted <0.05 in temporal analysis) encoding ligands of ECs in both cocultured and monocultured condition. Likelihood ratio test are performed for all the graphs. *P < 0.05, **P < 0.01, ***P < 0.001, and ****P < 0.0001 across all condition.

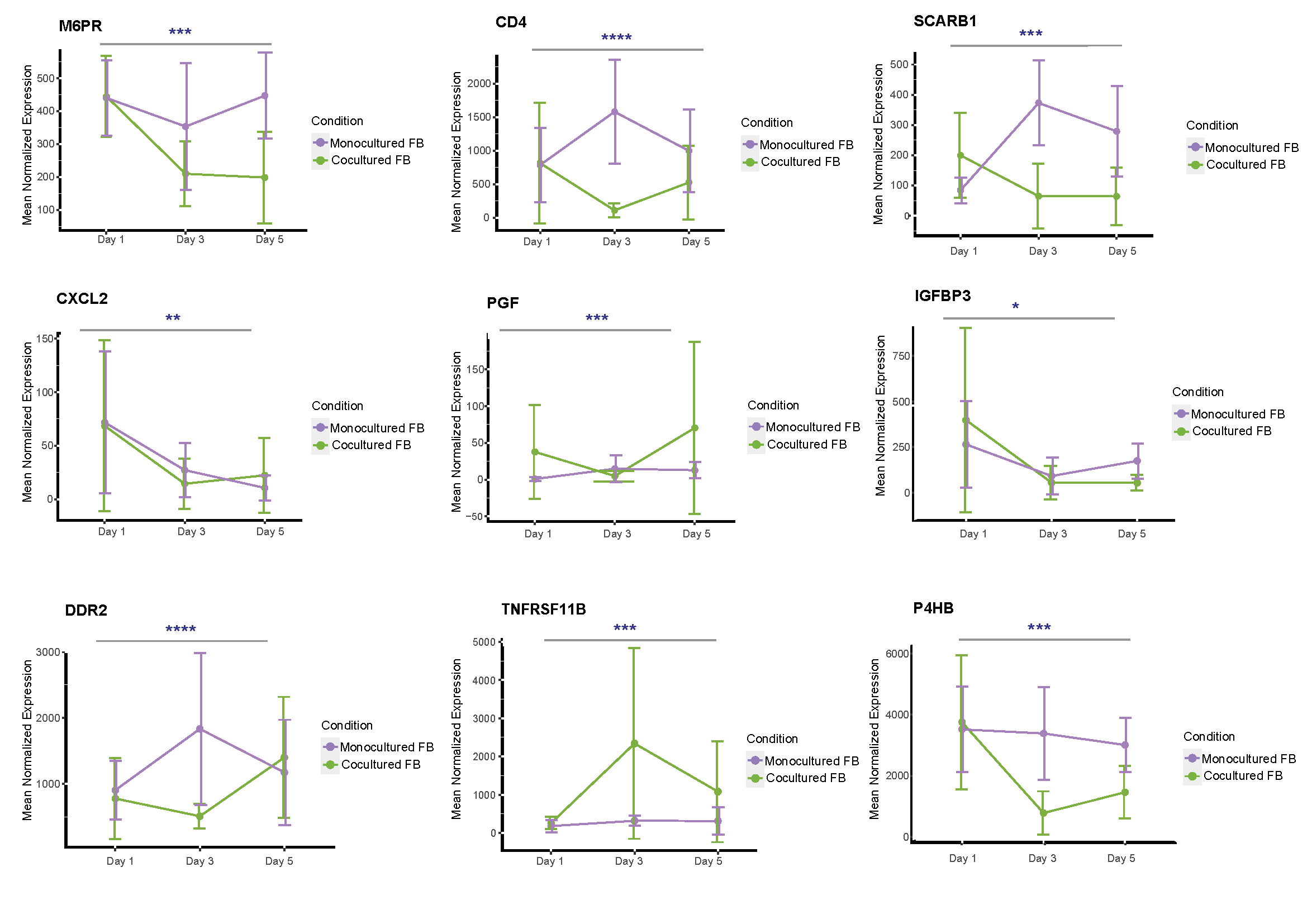

**Fig. S7. Temporal gene expression patterns of FB genes encoding receptor or ligands.**
Line plots showing the trend of DEGs (P-adjusted <0.05 in temporal analysis) encoding ligands of FBs in both cocultured and monocultured condition. Likelihood ratio test are performed for all the graphs. *P < 0.05, **P < 0.01, ***P < 0.001, and ****P < 0.0001 across all condition.

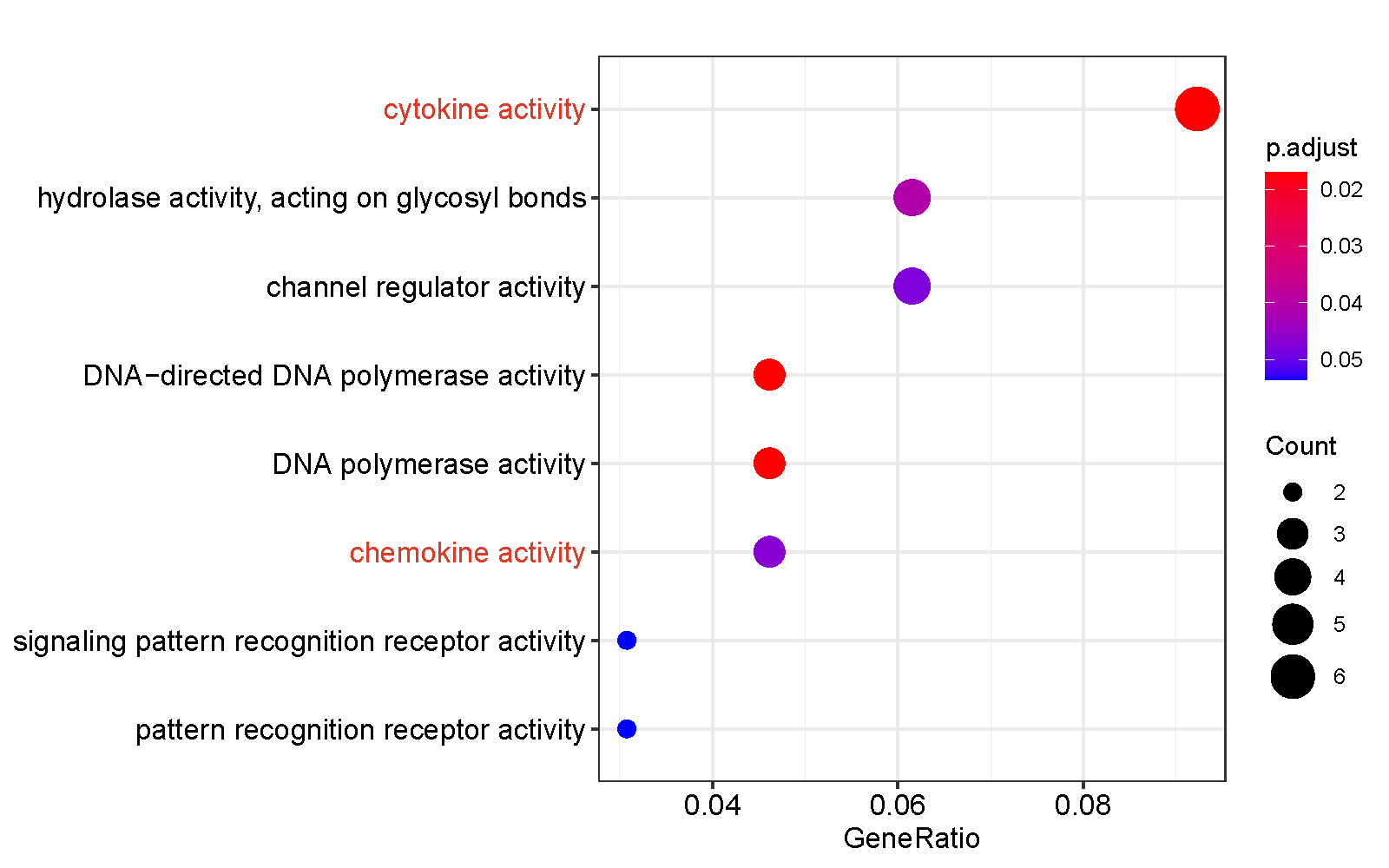

**Fig. S8. GO analysis of cluster 4 genes** **showing transcriptomic changes for coculture versus monoculture cells across three time points.**
The group of genes with higher expression at both day 1 and day 3 are shown to be associated with cytokines and chemokines related pathways (highlighted in red).

**Table S1. A full list of genes found to be genes to be differentially expressed between the monocultured and cocultured ECs (P adjusted <0.05, log2 fold change >1).**

(Separate file)
